# Age-dependent regulation of axoglial interactions and behavior by oligodendrocyte AnkyrinG

**DOI:** 10.1101/2024.04.01.587609

**Authors:** Xiaoyun Ding, Yu Wu, Victoria Rodriguez, Emily Ricco, James T. Okoh, Yanhong Liu, Daniel C. Kraushaar, Matthew N. Rasband

**Affiliations:** Department of Neuroscience, Baylor College of Medicine, Houston, TX 77030; Genomic and RNA Profiling Core, Baylor College of Medicine, Houston, TX 77030

## Abstract

The bipolar disorder (BD) risk gene *ANK3* encodes the scaffolding protein AnkyrinG (AnkG). In neurons, AnkG regulates polarity and ion channel clustering at axon initial segments and nodes of Ranvier. Disruption of neuronal AnkG causes BD-like phenotypes in mice. During development, AnkG is also expressed at comparable levels in oligodendrocytes and facilitates the efficient assembly of paranodal junctions. However, the physiological roles of glial AnkG in the mature nervous system, and its contributions to BD-like phenotypes, remain unexplored. Here, we generated oligodendroglia-specific AnkG conditional knockout mice and observed the destabilization of axoglial interactions in aged but not young adult mice. In addition, these mice exhibited profound histological, electrophysiological, and behavioral pathophysiologies. Unbiased translatomic profiling revealed potential compensatory machineries. These results highlight the critical functions of glial AnkG in maintaining proper axoglial interactions throughout aging and suggests a previously unrecognized contribution of oligodendroglial AnkG to neuropsychiatric disorders.

## Introduction

The cytoskeletal scaffolding protein AnkyrinG (AnkG) is a master regulator of ion channel clustering and neuronal polarity ^1,2^. In neurons, the ankyrin-repeat domain of AnkG clusters ion channels, transporters, receptors, and cell adhesion molecules to distinct membrane domains of the axon, while the spectrin-binding domain anchors them to the underlying periodic spectrin-actin cytoskeleton ^3^. Thus, AnkG dictates the formation of the axon initial segments (AIS) and nodes of Ranvier, where action potentials are initiated and efficiently propagated via saltatory conduction ^4^. The nodes of Ranvier are flanked by the paranodal junctions (PNJs) which are specialized axoglial junctions between myelinating oligodendrocytes and the axon. PNJs are essential for the formation and maintenance of the nodes of Ranvier and proper action potential conduction ^5,6^. As diffusion barriers, PNJs and their associated cytoskeleton restrict sodium channels to the nodes and voltage-gated potassium channels (Kv1) to the juxtaparanodes ^7,8^.

PNJs consist of the axonal cell adhesion molecules (CAMs) Contactin and Contactin-Associated Protein (Caspr), and the glial CAM Neurofascin (the 155 kDa isoform) (NF155) ^9–11^. The axonal CAMs recruit βII spectrin and protein 4.1b to create a cytoskeletal barrier that restricts ion channels and other membrane proteins to nodes and juxtaparanodes ^7,12,13^. In glia, NF155 is secured to the underlying cytoskeleton network by ankyrin scaffolding proteins—AnkG in the central nervous system (CNS) and AnkB in the peripheral nervous system (PNS). During early CNS development, glial AnkG facilitates the rapid and efficient assembly of paranodal junctions ^14^. However, the role of glial AnkG beyond early development remains unclear.

The AnkG coding gene, *ANK3*, is a risk gene for many neuropsychiatric disorders, including autism spectrum disorder (ASD), schizophrenia (SCZ), and bipolar disorder (BD) ^15–17^. Several genetic variants of *ANK3* have been identified by GWAS and a few validated using *Drosophila* or mouse models ^18^. For instance, *ANK3* heterozygous knockout mice demonstrate various behavioral abnormalities mimicking BD in humans ^19^, including decreased anxiety and increased motivation for reward. Other studies examined the underlying mechanism of BD-like phenotypes and contributions from different subtypes of neurons. For example, deletion of *ANK3* from either forebrain pyramidal neurons or PV interneurons ^20,21^ could each lead to robust phenotypes. These observations suggest that the underlying functions of *ANK3* and its association with BD risk may be multifaceted with contributions from many different cell types.

Despite a comparable level of *ANK3* transcripts in both neurons and oligodendrocytes ^22^, the functions of glial AnkG remain poorly understood, especially in adults. Therefore, we generated adult oligodendrocyte specific *ANK3* knockout mice and performed histological, physiological, and behavioral analyses as well as molecular and biochemical characterization of the altered oligodendrocytes. Our results reveal previously unknown functions of paranodal AnkG in the maintenance of the PNJ and suggest potential contributions of glial AnkG to psychiatric disease.

## Results

### Loss of oligodendroglial AnkG exacerbates age-related paranode disruption

Glial AnkG facilitates the rapid and efficient assembly of PNJs during early development^14^. However, whether glial AnkG is functionally required beyond the perinatal stage remains unknown. Therefore, we generated oligodendroglial specific conditional knock-out mice by crossing *Ank3^F/F^* with *NG2^Cre^* mice (AnkG cKO). Upon Cre-mediated recombination, exons 23 and 24 of *Ank3* are excised, resulting in premature stop codons in exons 25 and 26 (Fig. 1**a**). To confirm the efficient loss of AnkG protein in cKO animals, we performed immunoblotting of whole brain homogenates from 2-month-old mice (Fig. 1**b**), and we observed a significant reduction of the glial isoforms of AnkG (190 kDa and 270 kDa) ^14^ from cKO mice compared to littermate controls (Fig. 1**b**, **c**). To further evaluate the cell-type specificity of AnkG loss in AnkG cKO mice, we performed immunofluorescence staining of the optic nerve in 2-month-old mice. We found consistent loss of paranodal (glial) AnkG, but nodal (neuronal) AnkG remained unaffected (Fig. 1**d**, **arrowheads**).

**Figure 1.**
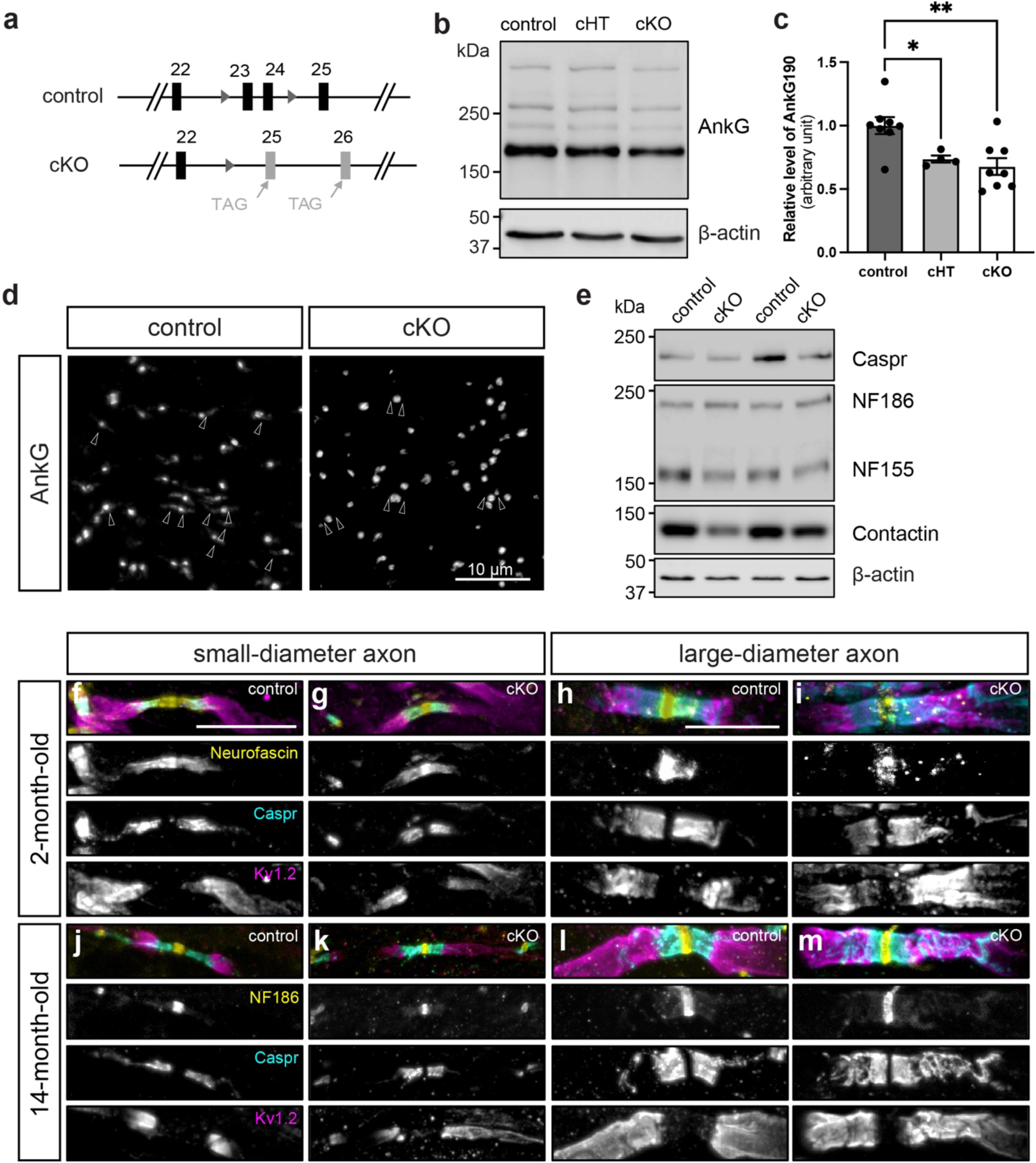
Loss of oligodendroglial AnkG leads to destabilization of paranodal junctions during aging. **a**, Schematic of the *Ank3* conditional allele with 2 loxp sites (triangles) flanking exons 23 and 24. Upon NG2^Cre^-mediated recombination, exons 23 and 24 are excised, introducing premature stop codons in exons 25 and 26. **b,** Immunoblot of whole brain homogenates from 2-month-old control (*Ank3^F/F^*), cHT (*NG2^Cre+^; Ank3^F/+^*), and cKO (*NG2^Cre+^; Ank3^F/F^*) animals. All three major isoforms of AnkG (480 kDa, 270 kDa, 190 kDa) are detected by guinea pig polyclonal AnkG antibody (SySy). β-actin serves as the loading control. **c,** Densitometry analysis of relative expression of AnkG (190 kDa isoform) in the brains of 2-month-old control, cHT, and cKO mice. Each data point represents an individual animal. One-way ANOVA with multiple comparison. *p=0.0436, **p=0.0031. **d,** Immunofluorescence staining of AnkG in the optic nerve of 2-month-old control and cKO mice. Arrowheads indicate paranodes. Scale bar, 10µm. **e,** Immunoblot of membrane-bound proteins from whole brain homogenates from 14-month-old control (*Ank3^F/F^*) and cKO (*NG2^Cre+^; Ank3^F/F^*) animals. Known PNJ components are detected by antibodies against Caspr, Neurofascin, and Contactin. β-actin serves as the loading control. **f**-**i,** Immunofluorescence labeling of nodal (Yellow: Neurofascin), paranodal (Yellow: Neurofascin, Cyan: Caspr) and juxtaparanodal (Magenta: Kv1.2) proteins on small-diameter axons (**f**, **g**) in the optic nerve of 2-month-old control mice, or on large-diameter axons in the spinal cord (**h**, **i**) of 2-month-old control mice. (**j**-**m**) Immunofluorescence labeling of nodal (Yellow: Neurofascin186), paranodal (Cyan: Caspr) and juxtaparanodal (Magenta: Kv1.2) proteins on small-diameter axons in the optic nerve of 14-month-old control mice (**j**) and cKO (**k**) mice. In the spinal cord of 14-month-old cKO mice, severe paranodal unwinding is only observed in the large-diameter axons (**l**, **m**) but not in the small-diameter axons.

To determine whether the molecular composition of the PNJs is altered in AnkG cKO mice, we extracted membrane-bound proteins from whole brain homogenates of AnkG cKO and control mice at 14 months of age. By immunoblotting with antibodies against known PNJ components, we detected a decrease in the levels of Caspr, Contactin, and glial Neurofascin (155 kDa). However, we saw no reduction in the neuronal Neurofascin isoform (186 kDa) in AnkG cKO mice (Fig. **1e**).

To determine if the PNJ structure is disrupted in young AnkG cKO mice, we performed immunofluorescence labeling for Neurofascin (nodes and paranodes), Caspr (paranodes), and Kv1.2 (juxtaparanodes; Fig. 1**f-i**). Consistent with our previous results using *CNP-Cre; Ank3^F/F^* mice ^14^, the PNJ structure was intact at 2 months of age in both the optic nerve (Fig. 1**f, g**) and spinal cord (Fig. 1**h, i**) of AnkG cKO mice. These data suggest that early developmental deletion of AnkG from oligodendrocytes may be effectively compensated for by redundant mechanisms in young adults.

Like AnkG cKO mice, we previously reported that loss of βII spectrin from Schwann cells also delayed the assembly of PNJ during development ^23^. Intriguingly, adult βII spectrin cKO mice were also phenotypically normal until 60 weeks of age, when they began to show disrupted paranodes in the PNS and CNS. Since AnkG and βII spectrin can interact—likely in a complex at the PNJ—we reasoned that loss of paranodal AnkG might also cause age-dependent deterioration of the PNJ. Therefore, we performed histological analyses of the paranodal structure in 14-month-old AnkG cKO animals (Fig. 1**j-m**) with gender-matched littermate controls. Interestingly, we observed no significant changes in the paranodal organization of small-diameter axons in the AnkG cKO optic nerve (Fig. 1**j, k**). However, there was clear disruption and “unwinding” of paranodal Caspr in large-diameter axons in the aged AnkG cKO spinal cord (Fig. 1**l, m**). These data suggest that oligodendroglial AnkG is required to maintain the PNJ in older adults.

### AnkG cKO mice show widespread histological changes

To determine the impact of PNJ disruption in older adults, we examined glial and neuronal cell markers throughout the CNS, including the optic nerve, brain, and spinal cord. For example, immunofluorescence staining of myelin basic protein (MBP) in the 14-month-old optic nerve showed ∼50% decrease in fluorescence intensity in AnkG cKO mice compared to their littermate controls (Fig. 2**a**, **b**), suggesting an overall reduction in myelin content in the optic nerve. We also observed a 2-fold increase in the fluorescence intensity of GFAP in the white matter of 14-month-old spinal cord, suggesting reactive astrogliosis (Fig. 2**a**, **c**). Proper myelination and axoglial interactions are crucial for neuronal function. Loss of myelin and disruption of PNJs is associated with axonal injury and neurodegeneration. One particularly susceptible type of neuron is the cerebellar Purkinje neuron. For example, both Caspr KO and NF155 KO mice show profound degeneration of Purkinje neuron axons ^24,25^. Thus, we further examined the effect of PNJ disassembly on Purkinje neurons in glial AnkG cKO mice. We observed a striking 50% loss of Purkinje neurons in 14-month-old AnkG cKO mice (Fig. 2**a**, **d**). Consistent with these histological observations, we found a reduction in MBP and Calbindin protein by immunoblot, and an increase in GFAP by immunoblot (Fig. 2**e**). Together, these results show that paranodal AnkG is critical for proper CNS myelination and Purkinje neuron survival, which may reflect disruption of PNJs.

**Figure 2.**
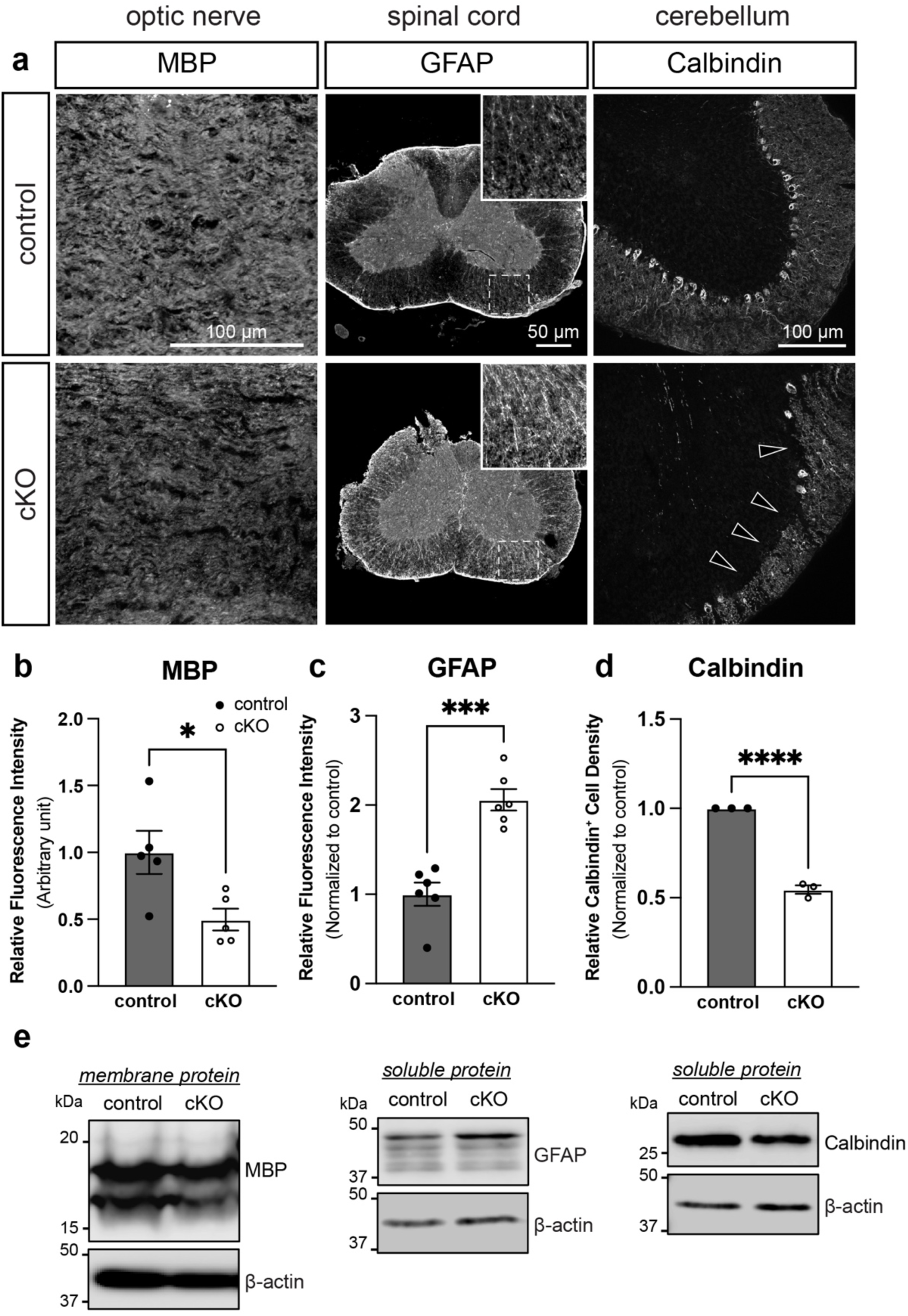
Paranodal destabilization in AnkG cKO mice leads to wide-spread CNS pathology. **a**, 14-month-old cKO mice show widespread histological changes in the entire CNS, including reduced myelin content in the optic nerve, increased reactive astrogliosis in the white matter of spinal cord, and Purkinje cell death (∼50%) in the cerebellum, indicated by the arrowheads. These phenotypes are consistent with paranodal disruption. **b**-**d,** Quantifications of fluorescence intensity of MBP and GFAP, and density of Calbindin+ cells. Each data point represents an individual animal. (unpaired t-test. p=0.0236, p=0.0001, p<0.0001.) **e,** Immunoblot analyses of whole brain homogenates confirm the reduction of MBP from the membrane-bound portion, as well as increase in GFAP and decrease of Calbindin from the soluble portion.

### AnkG cKO mice have slower action potential conduction

Proper myelination and axoglial interactions ensure efficient propagation of action potentials (APs). Given the disrupted paranodal structure and reduction in the MBP intensity observed in the optic nerve, we reasoned that the conduction velocity of action potentials in the CNS might be affected in AnkG cKO mice during aging. To evaluate AP conduction, we measured compound action potentials (CAPs) from the optic nerves of 12–16-month-old control and AnkG cKO mice using suction electrodes ^26^. CAPs were fitted with three gaussian curves corresponding to the three major groups of axons—slow, medium, and fast conducting axons in the optic nerve ^27^. We observed a significant broadening of the CAP (Fig. 3**a**), a decrease in maximum conduction velocity (Fig. 3**b**), and reduced conduction velocity for each subgroup of axons (Fig. 3**c**). Since disrupted paranodal junctions due to loss of paranodal CAMs are sufficient to reduce conduction velocities ^5,25^, we conclude the changes in CAPs are consistent with disrupted PNJs in aged AnkG cKO mice.

**Figure 3.**
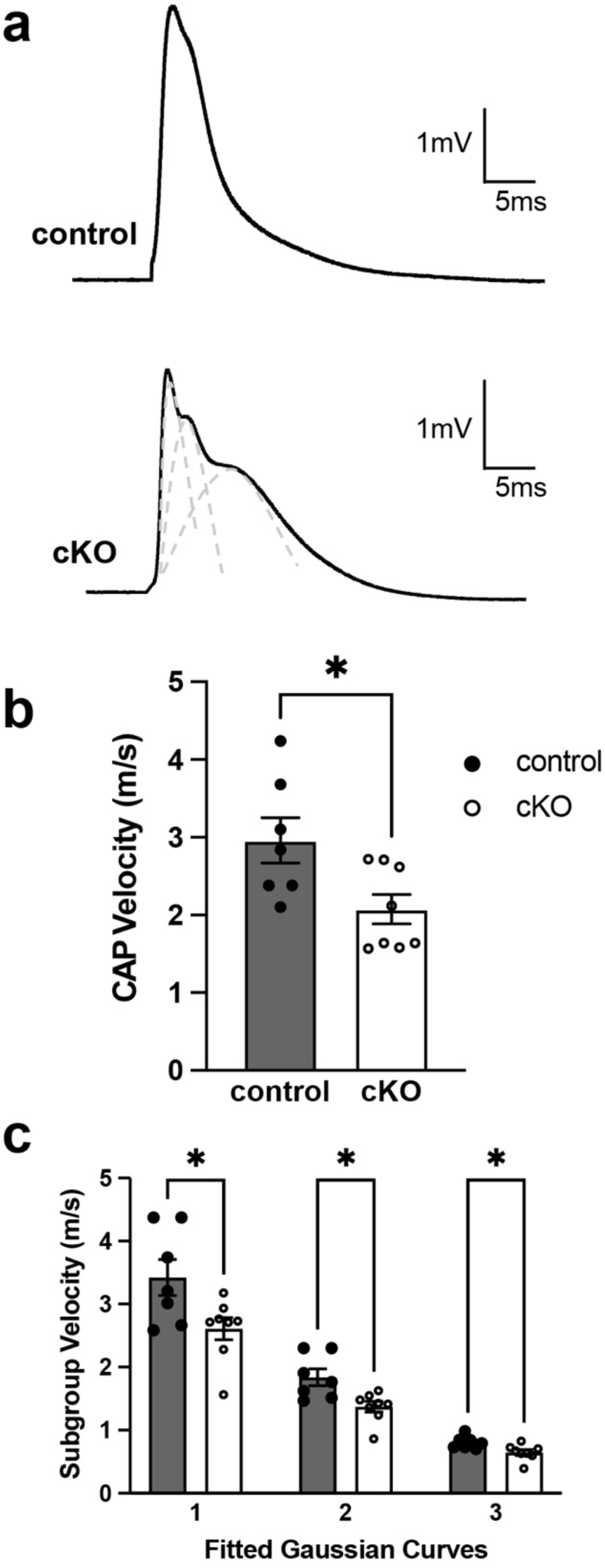
Loss of oligodendroglial AnkG results in slower conduction velocity. **a**, Representative compound action potential (CAP) traces recorded from freshly dissected 14-month-old optic nerves with suction electrodes ex vivo. **b,** The overall CAP velocity is calculated using the following formula: 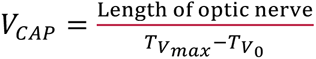 . The CAP velocity is significantly lower in AnkG cKO mice. Each data point represents an individual optic nerve. (Mann-Whitney test. p=0.0368.) **c,** Three Gaussian curves are fitted under each trace to differentiate fast-, medium- and slow-conducting axons. cKO mice show a slower conduction velocity in all three groups of axons. Each data point represents an individual optic nerve. (2-way ANOVA. p=0.0349, 0.0126, 0.0184 for subgroups 1, 2, and 3, respectively.)

### Loss of AnkG from oligodendrocytes reduces survival and overall health

The histological and physiological alterations in aged AnkG cKO mice suggested that the overall health and behavior of AnkG cKO mice could be impaired. To evaluate the overall health of the animals, we first examined postnatal survival rate. The Chi-square test showed a significant deviation of the AnkG cKO mice that are alive at P21 from the expected value based on the Mendelian ratio (Fig. 4**a**). Furthermore, both male and female AnkG cKO mice are significantly smaller in size and body weight compared to littermate controls starting from 20 weeks of age (Fig. 4**b**, **c**). As a result, AnkG cKO mice exhibited a shortened stride, stance, and sway in their gait patterns compared to littermate controls (Fig. 4**d**-**g**).

**Figure 4.**
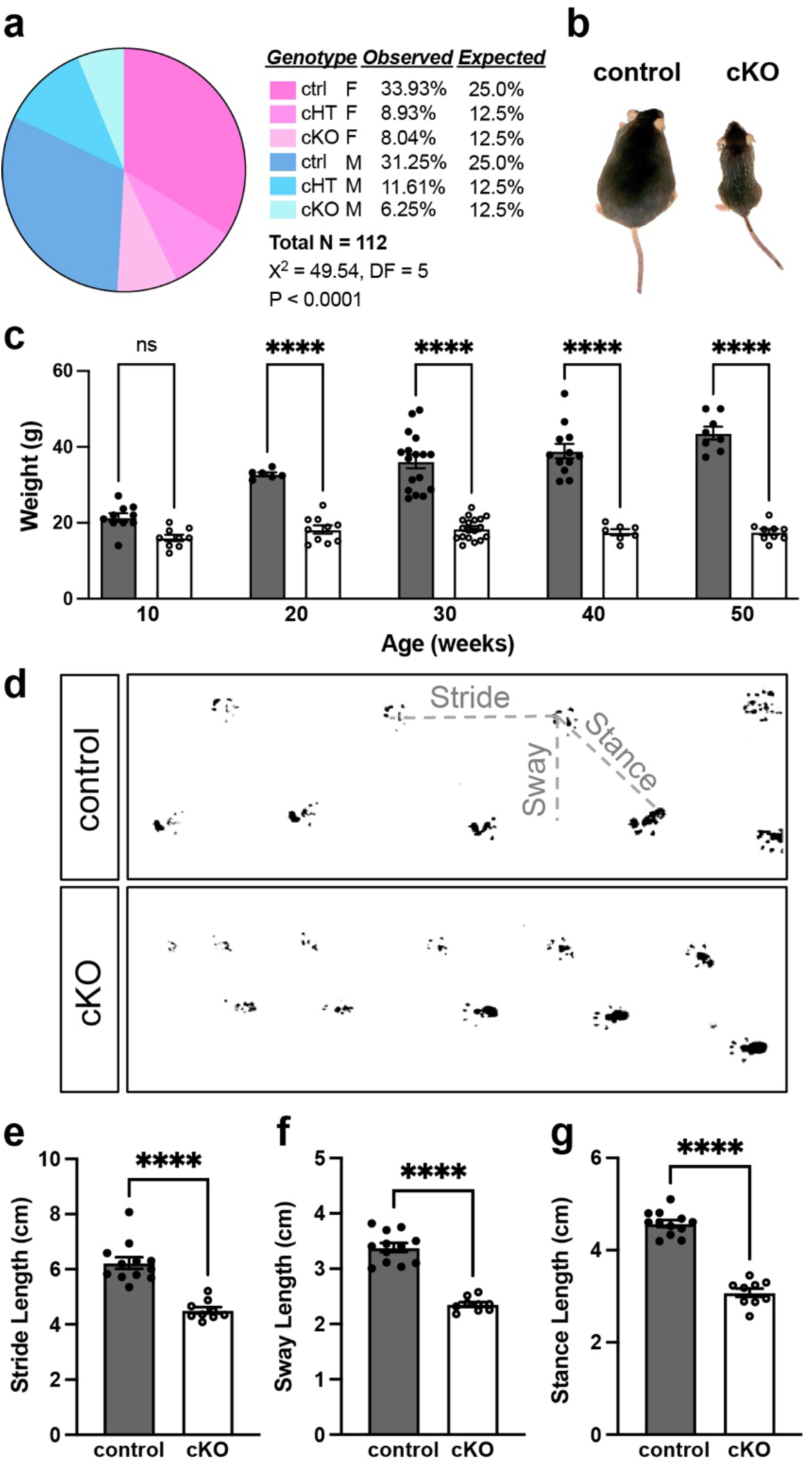
Loss of oligodendroglial AnkG is detrimental to survival and overall health. **a**, cKO mice show postnatal lethality with incomplete penetrance. Chi-square test. **b,** representative image of a 14-month-old AnkG cKO with its littermate control mice.**c,** Quantification of body weight at different ages. Each data point represents an individual animal. (2-way ANOVA. 10 weeks: p=0.06; for all other ages ****p<0.0001.) **d,** Representative image of hind-foot prints for gait analysis. **e**-**g,** Quantification of stride (**e**), sway (**f**), and stance (**g**) show significant reduction in all categories. Each data point represents an individual animal. (unpaired t-test. ****p<0.0001.)

### AnkG cKO mice have reduced locomotion and impaired motor learning

Since AnkG cKO mice lose Purkinje neurons at 14 months of age, we investigated their motor function. We performed the open field test in both young (2-4 months of age) and aged mice (> 9 months-old) and observed no differences in the total distance travelled by young cKO mice compared to control (Fig. 5**a**-**b**). However, the time young adult cKO mice spent in the inner zone was significantly decreased (Fig. 5**c**). There was a significant decrease in locomotor activity in aged cKO mice compared to control (Fig. 5**d**-**e**), without affecting the time spent in the inner zone. Thus, aged AnkG cKO mice have impaired locomotion.

**Figure 5.**
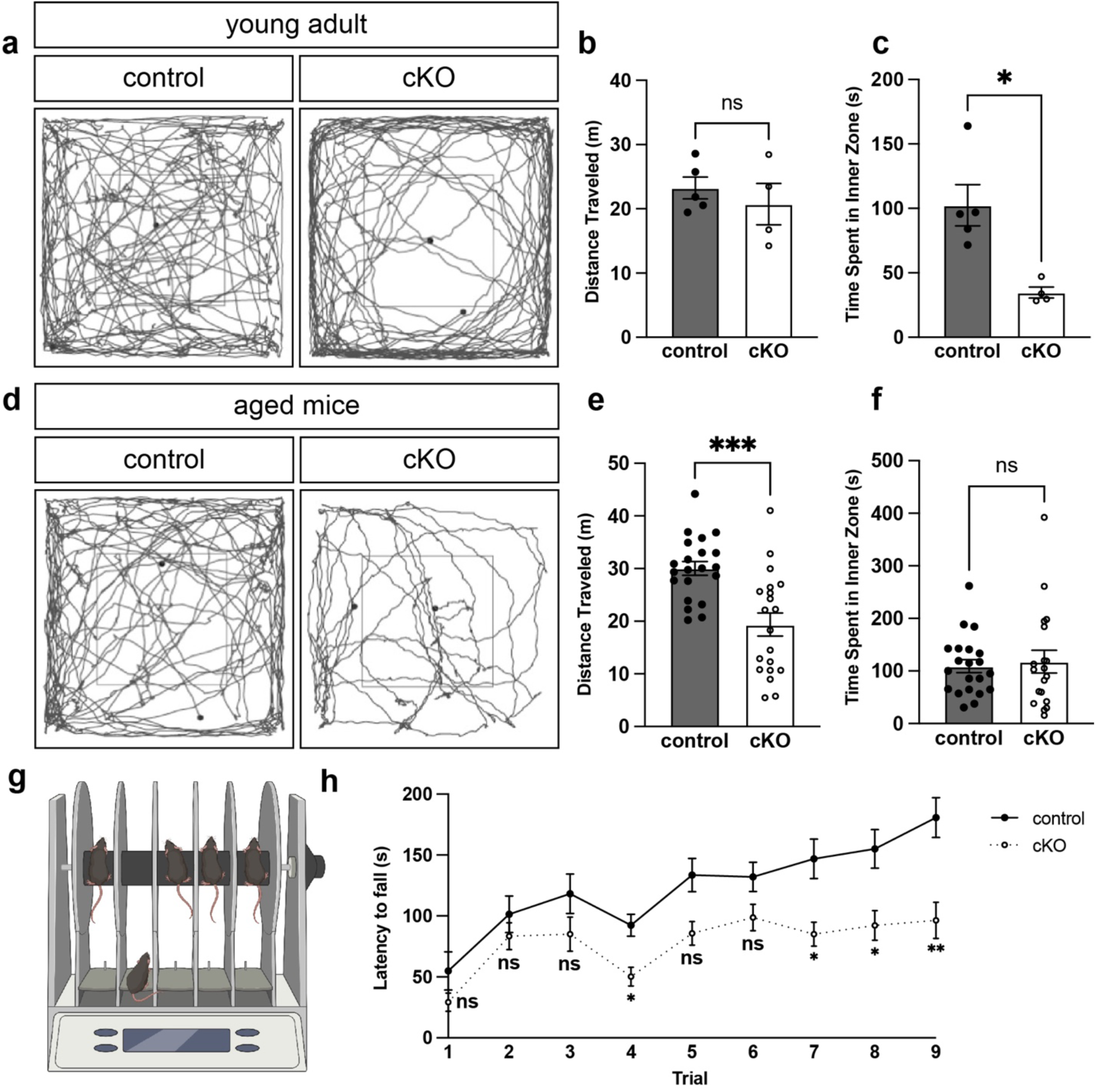
Loss of oligodendroglial AnkG leads to reduced locomotion and motor learning. **a**, Representative traces during 10-minute open field tests in 2 month-old control and AnkG cKO mice. **b,** Quantification of total distance traveled by each animal. Each data point represents an individual animal (unpaired t-test. p=0.4893). **c,** Quantification of time spent in the inner zone. Each data point represents an individual animal (Mann-Whitney test, p=0.0159). **d,** Representative traces during 10-minute open field tests in aged control and AnkG cKO mice (older than 6 months). **e,** Quantification of total distance traveled by each animal. Each data point represents an individual animal (unpaired t-test. p=0.0001). **f,** Quantification of time spent in the inner zone. Each data point represents an individual animal (Mann-Whitney test, p=0.7630). **g**-**h,** Accelerating rotarod test in 2 month-old control and AnkG cKO animals. Three trials per day were performed on three consecutive days. Data are presented as mean ± SEM. N=17 for control, N=13 for AnkG cKO. (2-way ANOVA. p=0.7794, 0.9780, 0.7277, 0.0123, 0.0728, 0.3681, 0.0281, 0.0358, 0.0063 for each consecutive trial, respectively).

Motor learning in young adults relies on active and adaptive myelination ^28^. To determine if motor learning is affected in young adult AnkG cKO mice (before Purkinje neuron degeneration), we performed the accelerating rotarod test for three consecutive days, with three trials each day (Fig. 5**g**). Over the course of training, control animals significantly extended their latency to fall, whereas AnkG cKO animals performed significantly worse than their littermate controls, with little improvement after training (Fig. 5**h**). Together, these results suggest that oligodendroglial AnkG is required for successful motor learning in young mice, before the onset of general motor function deterioration.

### AnkG cKO mice have altered social activity, anxiety, and depression-like behaviors

Myelin abnormalities and reduced white matter volume have been reported in patients with BD and SCZ ^29,30^. To determine whether loss of oligodendroglial AnkG is sufficient to cause behavioral abnormalities, we performed a battery of tests examining anxiety, depression-like behaviors, sociability, learning and memory, as well as SCZ-like behaviors. Using an elevated plus maze, we examined anxiety of AnkG cKO versus littermate control mice (Fig. 6**a**-**b**). Strikingly, AnkG cKO mice spent significantly more time exploring the open arm compared to control mice, suggesting that loss of glial AnkG has a strong anxiolytic effect, similar to the mania-like behaviors observed in previous studies of mice with altered AnkG expression in neurons ^19^, and impaired threat/stress signal processing in humans with *ANK3* variants ^31^.

**Figure 6.**
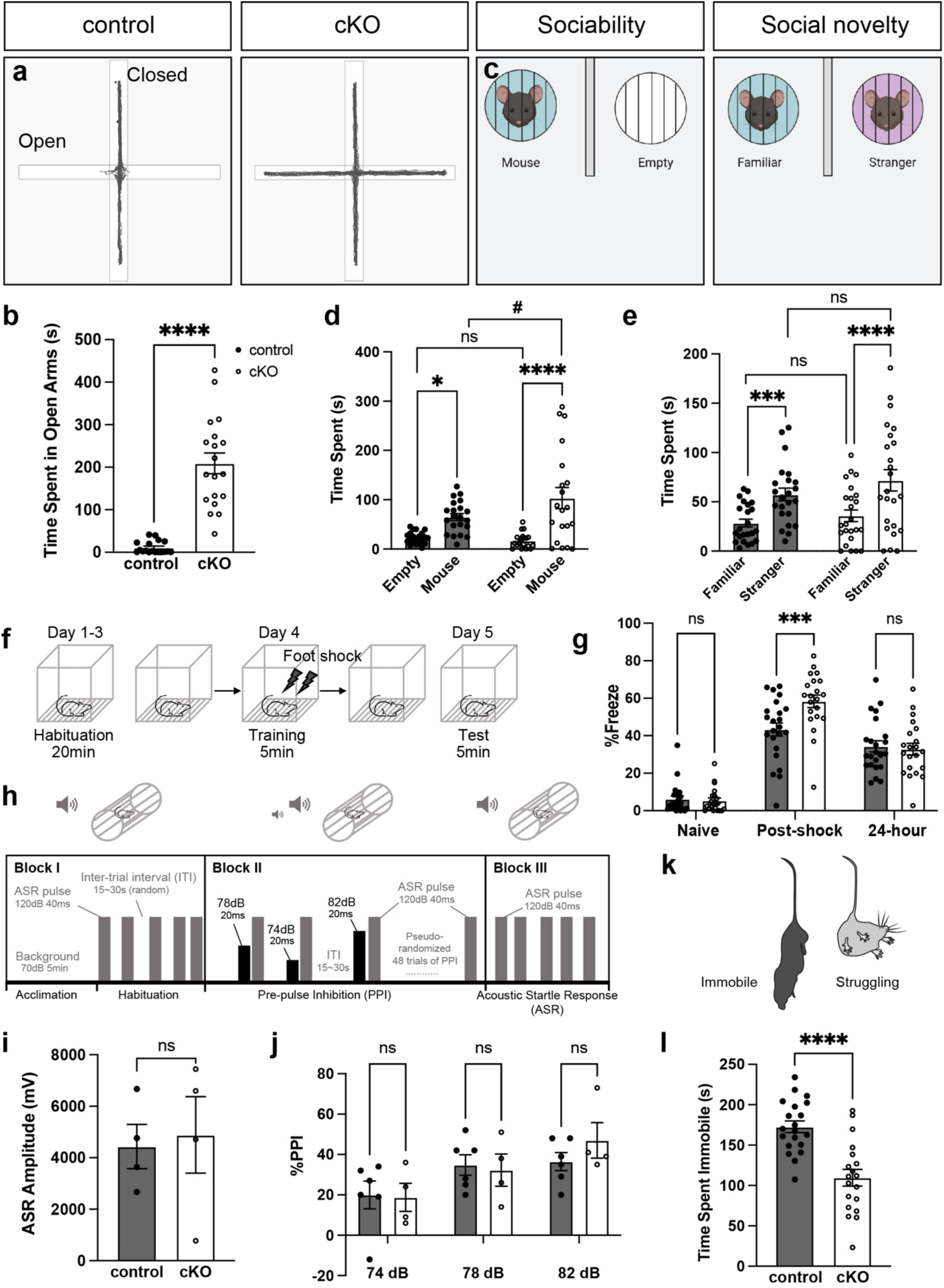
Loss of oligodendroglial AnkG leads to altered sociability, anxiety, and depression-like behaviors. **a**, Representative trace during 10-minute elevated plus maze test. **b,** Quantification of the time spent in open arms for control and AnkG cKO mice during the elevated plus maze test. (Mann-Whitney U test. p<0.0001.) **c,** Illustration of three-chamber social tests in control and AnkG cKO mice. **d,** Sociability measured by time spent with mouse versus empty cage. (2-way ANOVA with multiple comparisons. p*=0.0109, p****<0.0001, p=0.7112, p^#^=0.0174.) **e,** Social novelty measured by time spent with stranger versus familiar mice. (2-way ANOVA with multiple comparisons. p***=0.0007, p****<0.0001, p=0.4731, p=0.1713). **f,** Illustration of contextual fear conditioning test for learning and memory. **g,** Percentage of time spent freezing. (2-way ANOVA with multiple comparisons. p=0.9960, ***p=0.0007, p=0.9767.) **h,** Illustration of acoustic startle response and pre-pulse inhibition test. **i,** Amplitude of the startle response to 120 dB stimuli from AnkG cKO and littermate control mice. (unpaired t-test. p=0.8025.) **j,** %PPI for AnkG cKO and littermate control mice. (2way ANOVA with multiple comparisons. p>0.9999, p=0.9998, p=0.8671.) **k,** Illustration of tail suspension test for depression-like behavior in AnkG cKO and littermate control mice cKO mice. **i,** Quantification of immobility during the tail suspension test. (unpaired t-test. p<0.0001).

To determine the sociability and response to social novelty of AnkG cKO mice, we used the three-chamber test (Fig. 6**c**). We found that AnkG cKO mice showed significantly higher preference to mice than empty cups, and this was also significantly higher than control mice (Fig. 6**d**). AnkG cKO mice also had a higher preference towards “stranger” mice compared to familiar mice (Fig. 6**e**), but to a degree similar to those of control mice. Thus, AnkG cKO mice have heightened sociability but a normal response to social novelty.

To determine whether AnkG cKO mice have learning and memory defects, we performed the contextual fear conditioning test (Fig. 6**f**). During the training phase, AnkG cKO mice exhibited a significantly higher percentage of freezing behavior immediately after the foot shock compared to control mice (Fig. 6**g**). However, their ability to recall the stressful events 24 hours post training remained similar to control mice (Fig. 6**g**). Together, these results suggest that AnkG cKO mice have heightened social activity, but normal learning and memory functions.

Besides being a risk gene for BD, *ANK3* has also been implicated as a risk factor for SCZ^32^. To examine whether AnkG cKO animals exhibit SCZ-like behaviors, we performed acoustic startle response (ASR) and pre-pulse inhibition (PPI) tests (Fig. 6**h**). The ASR tests for the fast, protective, reflex response to a sudden loud noise; this test is a useful clinical tool in diagnosing patients. PPI further examines the sensorimotor gating mechanism. In a healthy subject, when a weak stimulus (pre-pulse) is given immediately prior to a strong startle stimulus, the reflex will be significantly suppressed. Impairment in PPI is often observed in patients with SCZ. Interestingly, AnkG cKO mice exhibited normal ASR (Fig. 6**i**) and normal PPI (Fig. 6**j**), suggesting the divergent role of glial *ANK3* in BD-like phenotypes and SCZ-related behavioral abnormalities.

Bipolar disorder in humans manifests as oscillating episodes of both mania and depression. To test whether AnkG cKO mice exhibit behavioral abnormalities related to depression, we performed the forced swim test (FST) and tail suspension test (TST). Surprisingly, AnkG cKO adults were unable to swim regardless of age, precluding the evaluation of depression-like behaviors using FST. However, during the TST test (Fig. 6**k**), AnkG cKO mice exhibited less immobile time compared to control mice, suggesting a lowered level of depression-like behavior in AnkG cKO mice (Fig. 6**l**).

### Inducible loss of oligodendroglial AnkG in young adult mice has mild effects

We previously showed that loss of AnkG delays the developmental assembly of paranodes ^14^. However, the effect of such a delay is limited to the perinatal stage. To further uncouple pathologies in the adult from disruption of paranodes during early development, we generated inducible oligodendroglial conditional *Ank3* knockout mice (*NG2^CreERTM^; Ank3^F/F^*, AnkG icKO mice). Tamoxifen was injected into 2-month-old adult mice, long after the PNJs were formed (Fig. 7**a**). After 1 month of recombination, we confirmed efficient and specific deletion of paranodal AnkG from the optic nerve (Fig. 7**b**-**c**). Despite the loss of paranodal AnkG (Fig. 7**b**-**c**, **arrowheads**), 3-month-old AnkG icKO mice had no defects in their PNJs in the optic nerve (Fig. 7**d**-**e**) since both Caspr and Kv1 channels were properly clustered at paranodes and juxtaparanodes, respectively.

**Figure 7.**
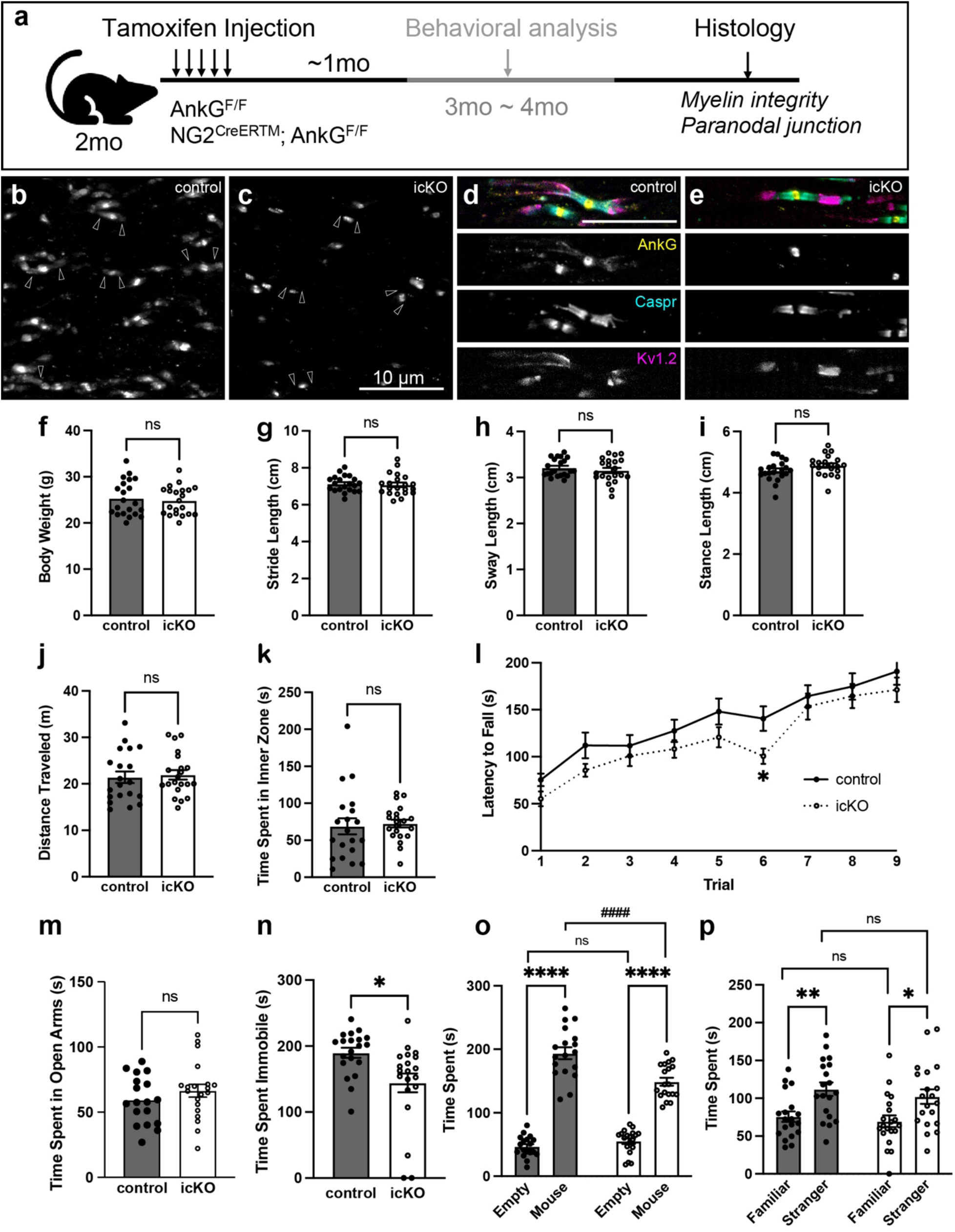
Loss of AnkG from adult mice. **a**, Illustration of inducible loss of AnkG from oligodendrocytes in NG2^CreERTM^; Ank3 ^F/F^ mice (AnkG icKO). **b**-**c,** Immunofluorescence staining of AnkG in the optic nerve of 3-month-old AnkG icKO mice (**b**) with littermate control (**c**). **d**-**e,** Immunofluorescence staining of the node, paranode, and juxtaparanode in the optic nerve of 3-month-old AnkG icKO mice (**d**) with littermate control (**e**) (Yellow: AnkG, Cyan: Caspr, Magenta: Kv1.2). **f,** Body weight of AnkG icKO vs control mice. (unpaired t-test, p=0.6588) **g-I,** Gait analyses of AnkG icKO vs control mice. (unpaired t-test for stride, p=0.7416; Mann-Whitney test for sway, p=0.3963; unpaired t-test for stance, p=0.1112.) **j**-**k,** Open field test of AnkG icKO vs control mice. (unpaired t-test for distance, p=0.7450; Mann-Whitney test for time in inner zone, p=0.3033.) **l,** Latency to fall from the accelerating rotarod for AnkG icKO vs control mice. (2-way ANOVA with multiple comparisons, p= 0.0576, 0.0935, 0.5004, 0.2071, 0.1259, 0.0141, 0.5468, 0.5935, 0.3180 for each trial, respectively). **m,** Time spent in open arms during the elevated plus maze test for AnkG icKO vs control mice. (Mann-Whitney test, p=0.3769.) **n,** Immobility during the suspension test for AnkG icKO vs control mice. (unpaired t-test, p=0.0417.) **o,** Three-chamber test of sociability for AnkG icKO vs control mice. (2-way ANOVA with multiple comparisons, p=0.3414, p****<0.0001, p^####^<0.0001.) **p,** Three-chamber test of social novelty test for AnkG icKO vs control mice. (2-way ANOVA with multiple comparisons, p=0.8263 for familiar, p=0.2136 for stranger, p**=0.0024, p*=0.0336.)

To further compare the AnkG icKO with AnkG cKO mice, we performed a battery of behavioral tests in AnkG icKO young adult mice. We found that loss of AnkG from young adult oligodendrocytes had minimal effects on health and behavior (Fig. 7**f**-**p**). Nevertheless, we did find a modest, but significant, reduction in depression-like behaviors seen in the tail suspension test (Fig. 7**n**). Since complex behaviors are orchestrated by many brain circuits that are already formed in young adults, our data suggest that the PNJ is likely more important during the development of these circuits rather than maintaining them in young adults. Thus, we conclude that disruption of glial AnkG during development has more severe and lasting effects than deleting it after development is complete.

### Translatomic profiling reveals compensatory machineries maintaining axoglial interactions in adults

In the absence of strong behavioral defects in the AnkG icKO mice, we sought to identify compensatory mechanisms that might contribute to preservation of the PNJ and oligodendrocyte function. Therefore, we performed unbiased translatomic profiling of oligodendrocytes from the whole brain homogenate of *NG2^CreERTM^;Ank3^F/F^;Rpl22^HA/+^* (AnkG icKO RiboTag) versus *NG2^CreERTM^;Rpl22^HA/+^* (control RiboTag) mice (Fig. 8**a**). This experiment reveals the mRNA transcripts actively engaged with ribosomes. Following tamoxifen injection and Cre-mediated recombination, we successfully immunoprecipitated ribosomes from NG2^+^ cells.

**Figure 8.**
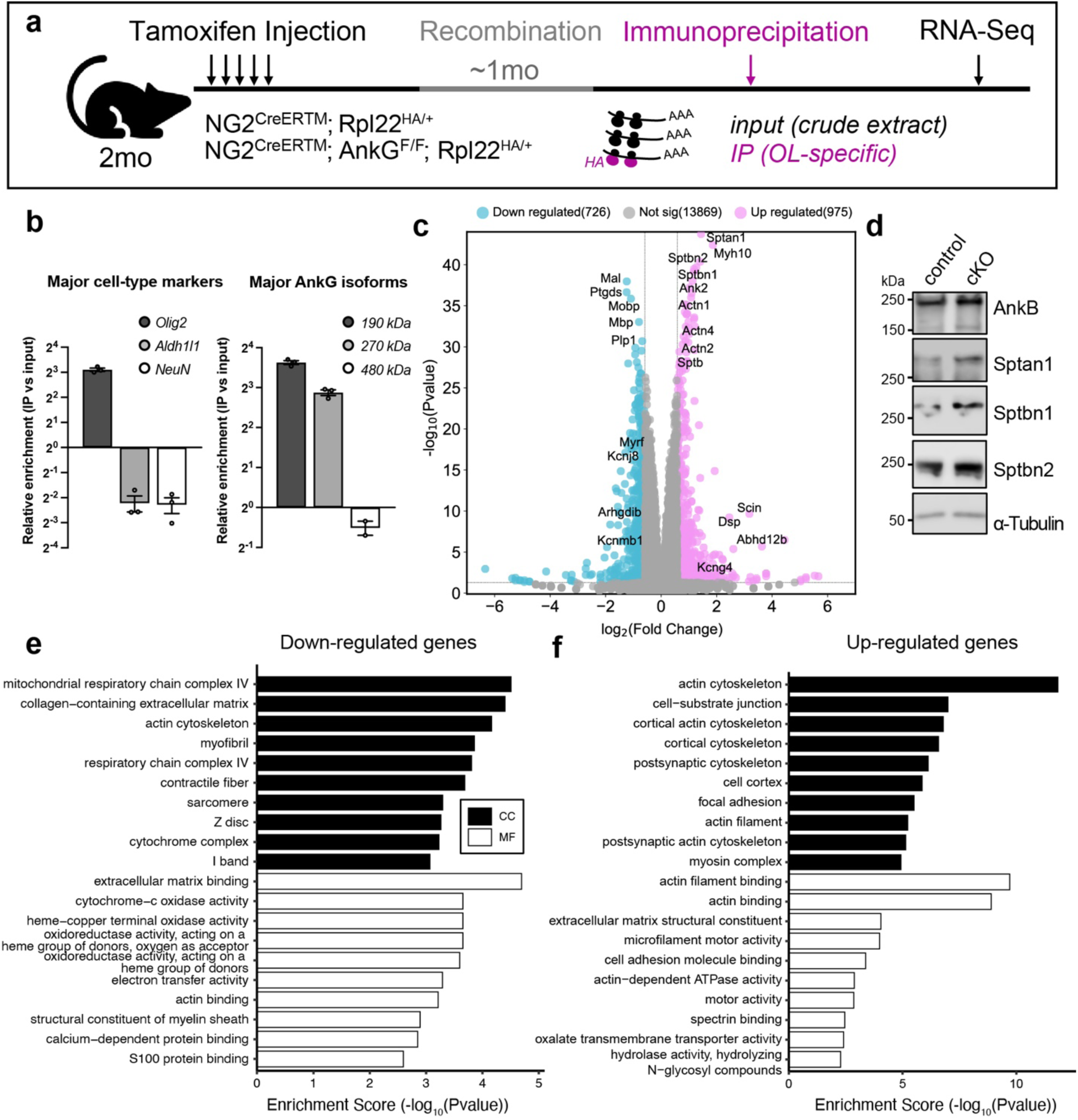
Translatomic profiling of AnkG-deficient oligodendrocytes reveals compensatory cytoskeletal mechanisms. **a**, Illustration of the experimental design for translatomic profiling in NG2^+^ oligodendroglial-lineage cells. **b,** Relative enrichment of major cell-type markers (oligodendrocytes: *Olig2*, astrocytes: *Aldh1l1*, neurons: *NeuN*) and major AnkG isoforms (190 kDa, 270 kDa, 480 kDa) by HA-IP. **c,** Volcano plot of differentially expressed genes between AnkG icKO IP sample and control IP sample. Magenta dots represent significantly upregulated genes, blue dots represent significantly downregulated genes, and gray dots represent genes that were not statistically differentially expressed (cutoffs: p < 0.05, |log_2_^FC^| > 0.6). The names of representative genes-of-interest are labeled in black. **d** Immunoblot of AnkB and spectrins to validate translatomic data. **e,** Bar plots for downregulated cellular component (CC, solid bar) and molecular functions (MF, empty bar) identified by Gene Ontology (GO) analyses. Enrichment scores are calculated by -log10(Pvalue). **f,** Bar plots for upregulated cellular component (CC, solid bar) and molecular functions (MF, empty bar) identified by Gene Ontology (GO) analyses. Enrichment scores are calculated by -log10(Pvalue).

Quantitative real-time PCR (qPCR) further confirmed ∼8-fold enrichment of the oligodendrocyte lineage-specific transcript *Olig2*, and ∼75% decrease of the astrocytic transcript *Aldh1l1* and neuronal transcript *NeuN* (Fig. 8**b**). Moreover, the oligodendroglial isoforms of AnkG (190 kDa and 270 kDa) were both over-represented in our immunoprecipitated (IP) samples compared to the crude extract (input), while the neuronal specific isoform (480 kDa) was significantly reduced after IP (Fig. 8**b**). These results are consistent with our previous study ^14^ and confirms the efficiency and specificity of the RiboTag system.

Among the differentially regulated transcripts (Fig. 8**c**, **blue**), we observed a general reduction of many myelin-related genes in AnkG icKO RiboTag mice compared to control RiboTag mice, including *Mal* (myelin and lymphocyte protein), *Ptgds* (prostaglandin D2 synthase), *Mobp* (myelin-associated oligodendrocyte basic protein), *Mbp* (myelin basic protein), *Plp1* (proteolipid protein 1), and *Myrf* (myelin regulatory factor) among others (Fig. 8**c** and Supplementary Data File 1). These translatomic changes were consistent with our observations in the histological and biochemical analyses using antibodies against MBP (Fig. 2**a**, **b**, **e**). These data strongly suggest that oligodendroglial AnkG is important for proper myelination of the CNS.

On the other hand, among the significantly upregulated genes (Fig. 8**c**, **magenta**), we also found a consistent increase of transcripts encoding the actin-spectrin-ankyrin cytoskeletal network, including *Sptan1* (α2 spectrin), *Sptb* (β1 spectrin), *Sptbn1* (β2 spectrin), *Sptbn2* (β3 spectrin), *Actn2* (α2 actinin), *Actn4* (α4 actinin), and *Ank2* (AnkB). To further validate these translational changes on the protein level, we performed immunoblotting using antibodies against some of the candidates, and we detected elevated level of α2, β1, β2 and β3 spectrins, as well as AnkB in the cKO whole brain homogenate compared to control (Fig. 8**d**). These data strongly suggest that compensation by AnkB and its binding partners, as well as the subsequent cytoskeletal remodeling, may ensure proper axoglial interactions in the absence of AnkG.

To further evaluate the overall impact of AnkG deletion from oligodendroglia, we then performed gene ontology analyses of down- and upregulated genes (Figs. **8e****, f**). Among all the downregulated cellular components (CC), we found a systematic reduction in mitochondrial respiratory chain, extracellular matrix, and actin cytoskeleton. Furthermore, extracellular matrix binding, oxidation, actin binding, and myelin sheath components were also significantly reduced among all molecular functions (MF; Fig. 8**e**). AnkG icKO RiboTag mice also had significant changes in opposition to the downregulated cellular components and molecular functions (Fig. 8**f**). For instance, the actin cytoskeleton, cell-substrate junction, focal adhesion, and myosin complex were all significantly enriched in the upregulated genes (Fig. 8**f**). As a result, the underlying molecular functions involving actin binding, extracellular matrix constituents, microfilament binding, cell adhesion molecule binding, spectrin binding, and actin-dependent ATPase and motor activity were also increased. That the actin cytoskeleton and extracellular matrix molecular functions are present in both the downregulated and upregulated pathways suggests these are redundant and compensatory molecular functions that maintain paranodal axoglial interactions after loss of glial AnkG. Overall, our unbiased translatomic profiling of AnkG icKO RiboTag mice strongly supports the conclusion that oligodendroglial AnkG regulates axoglial interactions at the paranode and identifies redundant and compensatory pathways that may also function to maintain axon-glia interactions.

## Discussion

Loss of neuronal AnkG contributes to bipolar-like behavioral abnormalities in mice ^19,21^. Since AnkG is the master regulator of neuronal polarity, and its functions at the AIS and nodes of Ranvier in neurons are well understood ^3,33^, it has been assumed that neuronal dysfunction underlies the BD-like behaviors in mice. However, AnkG has a comparable expression level in oligodendrocytes, but its roles in these glial cells are much less understood. Thus, we hypothesized that glial AnkG is also crucial for animal behavior. Here, we addressed three important questions: 1) is glial AnkG important for neuron-glial interactions, 2) is disruption of glial AnkG sufficient to alter the physiology and behaviors of mice, and 3) what molecular pathways are altered in the absence of glial AnkG? By generating oligodendroglial specific AnkG knockout animals, we explored the histological, physiological, behavioral, molecular, and biochemical features of these mice, and we showed the importance of oligodendroglial AnkG in aged animals. This study further highlights the pleotropic effects of AnkG in different cell types, at different developmental stages, as well as the potential contribution of glial *ANK3* to neuropsychiatric disorders like BD. Previous studies reported the existence of *ANK3* isoforms as risk factors for bipolar disorder ^34,35^. Future studies to investigate these specific isoforms may be very instructive and provide additional insight into the potential contributions of *ANK3* to BD.

The PNJs are the largest junctional adhesion complexes in vertebrates. Maintaining their structural integrity is pivotal to an efficient and functional nervous system across all myelinated organisms. Nodes of Ranvier are assembled through two overlapping glial mechanisms that converge on the axonal cytoskeleton ^4^. In the CNS, the axoglial-dependent assembly of the PNJ cytoskeleton is the primary node assembly mechanism, with a second mechanism that depends on a complex set of extracellular matrix molecules that can stabilize and retain nodal cell adhesion molecules for subsequent binding to axonal scaffolds and ion channels ^26^. The intricate and overlapping systems for node assembly highlights the evolutionary significance of redundant regulatory mechanisms. Our study focused on the maintenance of axoglial junctions at the paranodes, which corroborates the same principle of increasing the safety factor to ensure an efficient and functional nervous system throughout lifespan. Besides the role of the scaffolding protein AnkG in maintaining the PNJs, our unbiased translatomic profiling in oligodendrocytes in the absence of AnkG at the PNJ reveals both systematic remodeling of the spectrin-actin cytoskeletal network and dramatic changes in extracellular matrix components.

During aging, many developmental processes are re-employed to repair or replace damaged cells, tissues, and organs. Meanwhile, it is also a naturally perturbed system, where the extracellular environment, intercellular interactions, and intracellular properties of the aged cells are distinct from their developing counterparts. Therefore, aged animals provide a unique opportunity to further dissect out a well-compensated and complex system like the PNJs. In younger adult animals, we found the compensatory mechanisms are overwhelmingly effective, which could potentially mask the critical functions of certain proteins of interest. In this study, we confirmed that in young animals, AnkB may compensate for the loss of AnkG ^14^, but such compensation is not sufficient to maintain the deteriorating PNJ during aging. Previous studies showed that in aged CNS, paranodal reorganization results in the loss of transverse bands ^36^. In AnkG cKO mice, such a process may be accelerated. In aged Caspr mutant mice, disruption of the axon-glia interactions causes axonal degeneration and neuronal cell death ^37^. Similarly, deletion of βII spectrin from glial cells causes weakened muscles and slower AP conduction in sciatic nerves in aged but not young adults ^23^ . Based on these previous studies, we speculate that the paranodal structure is stable throughout early life due to compensatory mechanisms, however, these mechanisms are not sufficient to maintain the PNJ in aged AnkG cKO mice, perhaps due to lower affinity for the paranodal CAMs than the normal paranodal cytoskeletal and scaffolding proteins.

Our translatomic data also show that AnkG might be important for the maturation of oligodendrocytes, as many markers of mature oligodendrocytes are significantly reduced in the absence of AnkG. Previous studies showed that AnkG regulates neurogenesis and Wnt signaling by altering the subcellular localization of β-catenin ^38^. Wnt signaling components are well-known modulators of the differentiation, maturation, and formation of myelinating oligodendrocytes from oligodendrocyte precursor cells. Therefore, it is possible that the reduction in mature oligodendrocyte markers is partly due to aberrant Wnt signaling activity. More experiments are needed to test this hypothesis.

Despite our careful study design, this work has some limitations. First, our analyses rely on the Cre-lox system, which has the potential risk of “leakiness” into non-targeted cells, as well as the potential inefficient recombination. Our reporter analyses of the NG2^Cre^ mice showed around 1-5% recombination in neurons (varying among different animals) in addition to oligodendrocytes (data not shown). Furthermore, NG2 is also expressed in pericytes, which could potentially contribute to the effects we observed in these cKO mice. On the other hand, despite higher lineage-specificity, the tamoxifen induced NG2^CreERTM^ mice have only 25-50% efficiency in deleting Ank3, depending on the brain region. Therefore, the dramatic differences between NG2^Cre^ and NG2^CreERTM^ mice could be due to the limitations of the current genetic tools. Furthermore, this study was not meant to create a mouse model that mimics human bipolar disorder, as it is a complex disease that mice do not develop. Therefore, our study only strongly suggests that glial cells may contribute to the abnormal behaviors shown in BD patients. Future experiments with better tools will be needed to definitively address these questions.

## Methods

### ANIMALS

Oligodendroglial lineage specific AnkG conditional knockout mice (AnkG cKO) were generated by crossing *Ank3^F/F^* mice with *NG2^Cre^* mice. This results in the deletion of exons 22 and 23 and a premature stop codon in exon 24. *Ank3^F/F^* mice (B6.129-Ank3^tm^^2.1Bnt^/J) were generated as previously described ^39^ (RRID:IMSR_JAX:029797). *NG2^Cre^* mice were obtained from The Jackson Laboratory (RRID:IMSR_JAX:008533). Only female *NG2^Cre^* mice were used as breeders to avoid potential germ-line transmission. Both female and male mice were used for experiments. AnkG inducible conditional knockout mice (AnkG icKO) were generated using *NG2^CreERTM^* mice (RRID:IMSR_JAX:008538). Tamoxifen (20 mg/ml in corn coil) was administered intraperitoneally for 5 consecutive days (100 mg/kg of body weight) to 8-week-old mice which were analyzed after one month. RiboTag mice (B6J.129(Cg)-Rpl22tm1.1Psam/SjJ) were obtained from The Jackson Laboratory (RRID:IMSR_JAX:029977)

Genotyping for all transgenic animals was carried out by standard PCR on tail clips and further confirmed by immunofluorescence and or immunoblotting. Both specificity and efficiency of the Cre lines were assessed using an Ai3 reporter line. The following primers were used for genotyping:

*Ank3^F/F^* Forward: 5’- TTA ATT TGG GGA GGG GGG AGT C - 3’ *Ank3^F/F^* Reverse: 5’- TTG GGA TGC TTT GAT TCA GGG - 3’ *Cre* Forward: 5’ - TGC TGT TTC ACT GGT TAT GCG G - 3’

*Cre* Reverse: 5’ - TTG CCC CTG TTT CAC TAT CCA G - 3’ *RiboTag* Forward: 5’ - GGG AGG CTT GCT GGA TAT G - 3’ *RiboTag* Reverse: 5’ - TTT CCA GAC ACA GGC TAA GTA CAC - 3’

Mice were kept on a 12 h/12 h light/dark cycle (lights on at 7:00 am) and had access to food and water *ad libitum*. All animal care and experimental procedures were conducted in compliance with the National Institutes of Health Guide for the Care and Use of Laboratory Animals and were approved by the Baylor College of Medicine Institutional Animal Care and Use Committee (IACUC: AN4634).

### BEHAVIORAL ANALYSES

Similar numbers of male and female littermate mice were used for behavioral experiments. The age and number of animals are specified for each experiment and indicated in the figure legend. Whenever no sex-specific differences were observed, male and female mice were pooled together based on age and genotypes for statistical analyses. To eliminate odor cues, all apparatuses were cleaned with ethanol, dried, and ventilated between individual mice during testing. Mice were handled by the experimenter for 5 consecutive days (15 minutes/day) before the series of behavioral experiments. Animals were tested at the same time each day during the light cycle with the experimenter blinded to the genotypes.

#### Gait analysis

Gait characteristics were measured using footprint analysis as previously described ^40^ . In short, the hind feet of adult mice were painted with washable, nontoxic paint before walking through a 45 cm custom-made tunnel. The footprints were collected by white paper fitted to the floor and measured with ImageJ. For each animal, at least 20 representative and legible steps were analyzed, and the average value was used for quantitative analyses. Stride was measured from one left footprint to the next, sway was measured by the horizontal width between left and right foot, and stance was measured by the diagonal distance between left and right feet.

#### Open field test

The open field test was used to assess general locomotion. Briefly, adult animals were placed in the center of an open Plexiglass arena (40 × 40 cm, height 20 cm) and allowed to explore freely for 10 minutes. A camera and the AnyMaze software were used to perform automatic tracking of the animals. The parameters reported here include distance travelled and time spent in the inner zone throughout the test period. The inner zone was defined as the center 20 × 20 cm area.

#### Rotarod

The rotarod test was used to assess motor function and motor learning. Young adult mice (2- to 3-month-old) were acclimated to walking on the rotating rod at a constant speed of 4 rpm for 1 minute. After a 30-minute break, they were placed onto the rotarod accelerating from 4 rpm to 40 rpm for 5 minutes. Three trials were performed on each day for three consecutive days, with a 1-hour break in between two trials. The latency to fall was triggered by the lever or manually recorded—if the animal passively clutched onto the rod for two full rotations, the time was recorded as latency to fall.

#### Elevated plus maze

The elevated plus maze (EPM) test was used to assess anxiety and was performed as previously described ^41^ . The apparatus consisted of four metal arms, with two open and two enclosed arms perpendicular to each other (30 × 5 cm) and raised 40 cm above the floor. Adult mice were placed at the intersection between open and closed arms, with their heads towards the open arm. Animals were automatically tracked by a camera and the AnyMaze system. Time spent in each of the arms was reported.

#### Three-chamber social test

The three-chamber test was performed to assess the sociability and social novelty of the animals as previously described ^42^ . In brief, the test consisted of three 10-minute stages, including habituation, sociability, and novelty stages. In the first 10 minutes, adult mice were allowed to freely explore a 60 × 40 × 23 cm Plexiglass arena with three equally sized, interconnected chambers (left, center, and right). The left and right chamber each contained an empty wire-cup. During the next 10 minutes, the mice were placed back in the center, and another age- and gender-matched stranger animal was introduced to a randomly chosen wire-cup. In the last 10 minutes, a second stranger mouse was introduced into the remaining empty cup. Time spent interacting with each cup or mouse was recorded using a camera and the automated AnyMaze software and manually scored to include only the active interactions (sniffing, crawling upon, etc.). Sociability was evaluated by preference to animal versus empty cup, and social novelty was examined by preference to stranger versus familiar mouse.

#### Fear conditioning test

The fear conditioning test was performed to assess the learning and memory function as previously described ^43^ . Briefly, adult mice were accustomed to the conditioning chamber 20 minutes/day for 3 days. On the training day, mice received a foot shock (0.35 mA, 1 s) after 2 minutes in the conditioning chamber and remained in the chamber for another 2 minutes before returning to their home cage. 24 hours after training, mice were returned to the conditioning chamber for 5 minutes without any foot shock and the percentage of time spent freezing was recorded and analyzed by the FreezeFrame and FreeView software.

#### Acoustic startle response (ASR) and pre-pulse inhibition (PPI)

The ASR and PPI were used to assess schizophrenia-like behaviors in adult mice. In short, the subjects were acclimated to the animal enclosure 15 minutes/day for three days prior to the experiment, with a constant background noise playing at 70 dB. On the test day, the subjects were first accustomed to the animal enclosure for 5 minutes with 70dB background noise, after which they were given three blocks of different stimuli (Fig. 6h), namely, habituation, PPI, and ASR. During habituation stage, mice were given five 40 ms startle pulses at 120 dB, with random inter-trial intervals (ITI) of 15 to 30 seconds. The PPI stage consisted of 48 pseudo-randomized trials, in each trial, one of four 20 ms pre-pulses (70, 74, 78, and 82 dB) were given 100 ms prior to the 40 ms startle pulses at 120 dB. Pre-pulses were randomly presented 12 times at each decibel. The ITIs were randomly assigned between 15 to 30 seconds. During the ASR stage, each subject received five 40 ms 120 dB stimuli, with random ITI between 15-30 s. % PPI = 100 x (startle amp _ASR_ – startle amp _PPI_) / startle amp _ASR_.

#### Tail suspension test

The tail suspension test was performed to assess depression-like behaviors in adult mice ^44^ . To prevent tail-climbing behaviors, the tail of a subject mouse was first passed through a small plastic cylinder prior to suspension. The end of the tail was adhered to an adhesive tape that sticks back onto itself, with 2-3 mm of tail remaining outside of the tape. The free end of the tape was then applied to a horizontal suspension bar so that the approximate distance between the mouse’s nose and the apparatus floor was 20 to 25 cm. Each test lasted 6 minutes. A camera and the AnyMaze software were used to record and score the immobility of each animal.

### ELECTROPHYSIOLOGY

Compound action potential (CAP) recording was performed as previously described ^14^. Briefly, optic nerves were dissected out and submerged in oxygenated Locke’s solution bath with 1 mg/ml glucose. Both ends of the optic nerve were drawn into the suction electrodes, with stimulating electrode at the anterior end and recording electrode at the posterior end. Increasing currents were applied until a supra-maximal threshold was reached, at which the CAP was subsequently recorded. The conduction velocity was calculated by dividing the length of the recorded optic nerve by the latency (from stimulus onset to maximal peak). The recording traces of CAP were fitted into three Gaussian curves using python, to further assess the heterogeneous groups of axons with fast, medium, and slow conducting velocities.

### IMMUNOFLUORESCENCE STAINING

Mice were anesthetized with isoflurane, followed by transcardial perfusion with ice-cold PBS and then 4% paraformaldehyde (PFA). Tissues of interest were dissected out and post-fixed in 4% PFA at 4°C. Optic nerves were post-fixed for 30 minutes, transferred to 20% sucrose in 0.1 M phosphate buffer (PB) overnight, embedded in Tissue-Tek OCT (Sakura Finetek 4583) and sectioned using Microtome HS 450 at an interval of 10 µm. Knockout and littermate control tissues were mounted onto the same 1% bovine gelatin-coated glass coverslips, air-dried at room temperature before staining.

Brain and spinal cord were post-fixed at 4°C overnight, dehydrated in 20% sucrose in 0.1 M PB and 30% sucrose in 0.1 M PB at 4°C overnight. Tissues were embedded in OCT compound and stored at -80°C before sectioning. Cryostat sections were made at 20 µm intervals and mounted onto Superfrost Plus slides (Fisher Scientific Cat# 1255015), with knockout and littermate control tissues on the same slide for comparison.

Coverslips or Superfrost Plus slides with tissues were first washed three times, 5 minutes each, with 0.1 M PB to remove residue OCT. They were then blocked with phosphate buffer with detergent and goat serum (PBTGS: 0.1 M PB, 10% normal goat serum, 0.3∼0.5% Triton X-100) for 1 hour at room temperature. Primary antibodies diluted in PBTGS were added and incubated in a humid chamber at 4°C overnight. After washing three times with PBTGS, 5 minutes each, secondary antibodies diluted in PBTGS were added and incubated at room temperature for 1 hour. Samples were then washed with PBTGS, 0.1 M PB, and 0.05 M PB three times each before mounted with VECTASHIELD HardSet Antifade Mounting Medium (Vector Laboratories Cat# H-1400-10) and imaged with a Zeiss Apotome.2 Microscope.

### PROTEIN EXTRACTION AND IMMUNOBLOTTING

Animals were euthanized with isoflurane and immediately perfused transcardially with ice-cold PBS. Brains were quickly dissected out, snap-frozen in liquid nitrogen and stored at - 80°C. Frozen tissues were homogenized in glass Dounce Homogenizer on ice with the following buffer: 50 mM Tris-HCl, 15 mM EGTA, 10 0mM NaCl, and 1 mM DTT, supplemented with protease inhibitor cocktail (PIC) and PMSF immediately before use. Crude homogenates were first centrifuged at 2000 rpm at 4°C for 10 minutes to remove cell debris. The supernatant was then centrifuged at maximum speed at 4°C for another 90 minutes. Soluble proteins were present in the supernatant, and membrane-bound proteins were retrieved from the precipitated fraction. The concentration of each protein sample was measured using BCA assay (Pierce Cat# A55864) before boiling in SDS sample buffer at 95°C for 10 minutes and centrifuged at 5000 rpm for 5 minutes.

20 µg of total protein was resolved on 6% or 12% polyacrylamide gels using electrophoresis (SDS-PAGE) and transferred onto 0.45 µm nitrocellulose membrane using Bio-Rad Trans-blot system. Ponceau S was briefly added to the membrane to evaluate the quality of transfer and washed away using TBST (TBS with 0.1% Tween-20). The membrane was blocked with Blotto solution (5% skim milk in TBST) for 1hr at room temperature (RT). Primary antibodies were diluted in Blotto solution and incubated with the membrane at 4°C overnight. The membrane was then washed with TBST for three times, 10 minutes each, and then incubated with HRP-conjugated secondary antibodies in Blotto solution for 1 hour at room temperature. The membrane was then washed with TBST three times before developing signals using SuperSignal West Pico PLUS (Pierce Cat# 34580) and imaging with the Licor odyssey Fc system. Densitometry measurements of immunoblots were performed in FIJI/ImageJ (RRID:SCR_003070).

### ANTIBODIES

The following primary antibodies were used for standard immunofluorescence: Mouse monoclonal antibodies against AnkG (clone N106/65, RRID:AB_2877525, 1:500), Caspr (Clone K65/35, RRID:AB_2877274, 1:1000), Kv1.2 (Clone K14/16, RRID:AB_2877295, 1:1000), Calbindin (UC Davis/NIH NeuroMab Facility Cat# L109/57, RRID:AB_2877197, 1:1000), and MBP (Clone SMI 94, BioLegend Cat# 836504, RRID:AB_2616694, 1:500). A rabbit monoclonal antibody against Neurofascin 186 (Clone D6G6O, Cell Signaling Technology Cat# 15034, RRID:AB_2773024, 1:500). Rabbit polyclonal antibodies against Caspr (Abcam Cat# ab34151, RRID:AB_869934, 1:500), MBP (Abcam Cat# ab40390, RRID:AB_1141521, 1:500), and Calbindin D-28K (Millipore Cat# AB1778, RRID:AB_2068336, 1:1000). Chicken polyclonal antibodies against GFAP (Abcam Cat# ab134436, RRID:AB_2818977, 1:2000) and Neurofascin (R&D Systems, catalog #AF3235, RRID:AB_10890736, 1:500). A guinea pig polyclonal antibody against AnkG (Synaptic Systems Cat# 386 004, RRID:AB_2725774, 1:500).

The following primary antibodies were used for immunoblotting: Rabbit polyclonal antibodies against Caspr (Abcam Cat# ab34151, RRID:AB_869934), Calbindin D-28K (Millipore Cat# AB1778, RRID:AB_2068336), MBP (Abcam Cat# ab40390, RRID:AB_1141521), AnkB (Matthew Rasband Lab, E1588), and β3 Spectrin (Novus Cat# NB110-58346, RRID:AB_877723). A guinea pig polyclonal antibody against AnkG (Synaptic Systems Cat# 386 004, RRID:AB_2725774). A chicken polyclonal antibody against Neurofascin (R&D Systems, catalog #AF3235, RRID:AB_10890736). A mouse monoclonal antibody against Contactin (UC Davis/NIH NeuroMab Facility Cat# K73/20, RRID:AB_2877311), α2 Spectrin (BioLegend Cat# 803201, RRID:AB_2564660), β2 Spectrin (BD Biosciences Cat# 612563, RRID:AB_399854, α-Tubulin (Sigma-Aldrich Cat# T6074, RRID:AB_477582), and β-Actin (Sigma-Aldrich Cat# A1978, RRID:AB_476692).

All secondary antibodies were purchased from Thermo Fisher Scientific or Jackson ImmunoResearch Laboratories. Isotype specific secondary antibodies were used whenever possible.

### IMAGE PROCESSING

Immunofluorescence images were observed with an AxioImager (Carl Zeiss) with the Apotome optical sectioning module and acquired with a digital camera (AxioCam, Carl Zeiss). Images from the same experiments were acquired and processed in batch using the ZEN software (Carl Zeiss) under identical parameters. For z-stack images, maximum intensity projection (MIP) function was applied. For better visualization of representative PNJs, images were taken at the magnification of 100x and cropped in Adobe Photoshop software, with each channel isolated and inverted. Relative intensity measurements of fluorescence images and densitometry analysis of immunoblots were performed in FIJI/ImageJ (RRID:SCR_003070). The density of Purkinje cells was calculated by the number of Calbindin^+^ cells divided by the length of the Purkinje layer. The relative fold-changes were calculated by normalizing knockout to littermate controls. At least 3 images from 3 different animals per genotype were included for each analysis.

### STATISTICAL ANLYSIS

Statistical analyses for immunofluorescence images, immunoblots, and behavioral assays were performed in GraphPad Prism10 (RRID:SCR_002798). No power analyses were performed to predetermine the sample sizes, but our sample sizes are comparable to those previously reported ^14^ . For histological analyses, at least three age- and gender-matched littermates were selected per genotype. If there were fewer than three pairs, mice from another litter with similar birth date were added for statistical analyses. For electrophysiological analyses, traces from 8 optic nerves per genotype were analyzed.

The proper statistical tests were carefully selected based on normality and consideration of multiple comparisons. Data were analyzed using Microsoft Excel, python, and GraphPad Prism. The following tests were employed where appropriate: unpaired Student’s t-test, Mann-Whitney U test, one-way ANOVA, two-way ANOVA with multiple comparisons, Chi-square test, and Kolmogorov–Smirnov test. All error bars are ± SEM unless otherwise indicated. Each data point represents an individual animal unless otherwise stated.

### RNA PURIFICATION AND SEQUENCING

Actively translated mRNAs from the oligodendrocyte lineage were isolated and prepared as previously described ^45^ . Briefly, NG2^CreERTM^; RiboTag mice were injected with tamoxifen (20 mg/ml in corn coil) for 5 consecutive days (100 mg/kg of body weight) at 2 months of age and harvested 1 month later. Mice were anesthetized with isoflurane and perfused with ice-cold PBS, and the whole brain homogenates were immediately processed for IP-enrichment of ribosomes from oligodendrocytes. Three animals were pooled together for one RNA-seq sample. 10% of the cleared homogenate was collected as input. IP was performed overnight with HA-antibody (Biolegend Cat# 901505; RRID:AB_2565023) and Pierce protein A/G Magnetic Beads (Thermo Scientific Cat# 88803). RNA was isolated from both input and IP samples using RNeasy Mini Kit (Qiagen Cat# 74104).

NanoDrop spectrophotometry (ThermoFisher Scientific Inc.) was used to assess RNA purity while RNA integrity was evaluated using a 2100 Bioanalyzer with RNA chips (Agilent Technologies, Santa Clara, CA). The Genomic and RNA Profiling Core prepared libraries with the Takara SMARTer® Stranded Total RNA-Seq v3 - Pico Input Mammalian kit (Takara, p/n 634485). Using the KAPA Library Quantification kit for Illumina platforms (KAPA Biosystems, Wilmington, MA), libraries were quantitated and pooled equimolarly. Sequencing of mRNA libraries was performed on an Illumina NovaSeq 6000 platform to around 90 million read pairs per sample.

### CDNA SYNTHESIS AND QUANTITATIVE PCR

cDNA was synthesized from both IP and input RNA samples as previously described ^46^. Briefly, IP and input RNA samples were reverse transcribed into cDNA using SuperScript IV VILO MasterMix, followed by genomic DNA removal with ezDNase Enzyme (Invitrogen Cat# 11766050). And quantitative PCR (qPCR) was performed using iTaq Universal SYBR Green Supermix (Bio-Rad Cat# 1725122) using the StepOnePlus Real-Time PCR System. Melting curve plots were examined to ensure specificity. The relative expression of genes-of-interest was quantified using the 2^-ΔΔCT^ method, with *Rplp0* serving as the internal control.

The following primers were used for the qPCR:

*Olig2* Forward: GAACCCCGAAAGGTGTGGAT

*Olig2* Reverse: GCCCCAGGGATGATCTAAGC

*Aldh1l1* Forward: ATGATCATCTCTCGGTTTGCTGA

*Aldh1l1* Reverse: CATCGGTCTTGTTGTATGTGTTG

*Rbfox3 (NeuN)* Forward: GCGGTCGTGTATCAGGATGG

*Rbfox3 (NeuN)* Reverse: CGATGCTGTAGGTTGCTGTG

*AnkG-190* Forward: CTTTGCCTCCCTAGCTTTAC

*AnkG-190* Reverse: TCTGTCCAACTAAGTCCCAG

*AnkG-270* Forward: GCCATGTCTCCAGATGTTG

*AnkG-270* Reverse: TCTGTCCAACTAAGTCCCAG

*AnkG-480* Forward: GAGGCACCGCCCTTAAA

*AnkG-480* Reverse: GCCAGCTCTGTCCAACTAA

*Rplp0* Forward: CAGGCGTCCTCGTTGGAG

*Rplp0* Reverse: ATCTGCTGCATCTGCTTGGAG

### BIOINFORMATICS ANALYSIS

The first 14 nucleotides from Read 2 FastQ files were trimmed to remove UMI, UMI linker, and Pico v3 SMART UMI Adapter sequences. Illumina adapters were trimmed from raw reads using Cutadapt v4.4 ^47^, removing the following adapter sequences: Read1: AGATCGGAAGAGCACACGTCTGAACTCCAGTCA, Read2: AGATCGGAAGAGCGTCGTGTAGGGAAAGAGTGT. Quality trimming and filtering was performed using Trimmomatic v0.39 ^48^ to trim the first 5 base calls, to trim read segments that fall below a quality score of 25 using SLIDINGWINDOW:4:25, and retain reads with a minimum length of 100bp. Reads were then aligned to UCSC mm10 using STAR v2.7.10b ^49^ with gene quantification generated using (quantMode GeneCounts).

Raw counts were filtered based on CPM, retaining genes with CPM > 1.5 for at least 2 samples. Normalization was performed with the R program edgeR ^50^, TMM normalization. Differential gene expression analysis was also performed using edgeR under a negative binomial generalized linear model with quasi-likelihood dispersion estimates, assuming an FDR < 0.1 as significant.

The volcano plot of differentially expressed genes and bar plot of Gene Ontology and KEGG pathway enrichment analyses were generated by https://www.bioinformatics.com.cn/en.

## Acknowledgments

This work was supported by grants from the National Institutes of Health (5R01MH121544 to M.N.R.) and the Dr. Miriam and Sheldon G. Adelson Medical Research Foundation (M.N.R.). This project was also supported in part by the Genomic and RNA Profiling Core at Baylor College of Medicine with funding from the NIH S10 grant (1S10OD023469).

## Author contributions

Conceptualization, methodology, data curation, validation, investigation, visualization, and writing—original draft and editing: X.D.; Investigation, data curation: Y.W., V.R., E.R.; Resources: J.T.O. and Y.L.; Data curation, analysis, resources, and writing—original draft: D.C.K.; Conceptualization, methodology, data curation, writing—editing, project administration, and funding acquisition: M.N.R.

